# Human Immunodeficiency Virus Status in Malnourished Children seen at Lagos

**DOI:** 10.1101/357608

**Authors:** EO Temiye, OF Adeniyi, IB Fajolu, AA Ogbenna, TA Ladapo, CI Esezobor, AO Akinsulie, CA Mabogunje

## Abstract

**Introduction:** Human immunodeficiency virus and protein energy malnutrition are still prevalent in Nigeria and the occurrence of the two conditions together confers a poor prognosis. The aim of this study was to determine the current categories of malnutrition amongst under 5 children in Lagos, document their HIV status and determine any peculiarities in the clinical features, haematological and some biochemical profile in these children.

**Methods:** The study was a prospective crossectional study conducted at the Paediatric department of the Lagos University Teaching Hospital and the Massey Street Children’s Hospital over a 6 month period. All the subjects had anthropometry, HIV testing, full blood count and serum proteins done. The factors associated with HIV status were determined with the logistic regression analysis.

**Results:** Two hundred and fourteen (214) malnourished children ≤5 years were recruited into the study and 25(11.7%) were HIV positive. One hundred and five (49.1%) of the participants had moderate malnutrition while 25.2% had severe forms of malnutrition. Fever, cough and diarrhea were the commonest symptoms. Severe wasting, oral thrush, dermatoses and splenomegaly were seen more commonly in the HIV positive subjects. The haematological indices were comparable in the two groups, however, the total protein was significantly higher in the HIV positive subjects compared to the negative group (p=0.042). Multivariate analysis showed that the total protein (p=0.001) and platelet count (p=0.016) could significantly predict the occurrence of HIV in the malnourished children

**Conclusion:** The presence of severe wasting, oral thrush, diarrhea, splenomegaly, thrombocytopenia and high total proteins in malnourished children should heighten the suspicion of possible underlying associated HIV infection. This study reinforces the recommendation by the World Health Organisation (WHO) that all malnourished children should have mandatory HIV screening.

## Introduction

The Human Immunodeficiency Virus (HIV) infection and its attendant problems remain a threat to global health, more so in the sub Saharan continent where malnutrition is also prevalent^1^ and the paediatric age group remain the most affected vulnerable group.^1^ In developing countries, malnutrition contributes significantly to 50% of childhood mortality either directly or indirectly.^2^ The mortality that occurs from the malnourished state is significantly related to the body growth parameters, namely the weight for age, weight for height and height for age. ^2,3^ Severely malnourished children with z scores of the weight for height less than 3-standard deviation from the mean values (-3SD) are at a 10 fold risk of a higher mortality compared to children with a z-score higher or equal to 1.^2,3,4^

Malnutrition and HIV infection are believed to be intertwined in a vicious cycle as HIV increases the vulnerability to malnutrition while the latter lowers immunity and enhances vulnerability to HIV transmission risk and disease progression in HIV infected children. ^5–7^The clinical features of severe malnutrition and HIV/AIDS are known to overlap in young children. This may affect the accurate diagnosis of presumptive HIV infection in resource poor settings where HIV testing facilities are still inadequate and the two conditions often coexist. Malnutrition, although mainly attributed to nutritional deficiency, may be multifactorial in origin and the malnourished state which is usually accompanied by some degree of immunocompromise predisposes to infections generally and this does not exclude the HIV infection. The World Health Organization(WHO) actually classifies wasting as a Stage 3 AIDS condition and thus distinguishing malnutrition from the former clinically sometimes can be difficult,^7–8^ thus, the recommendations remains that all malnourished children should have HIV testing done.^8^

The prevalence of HIV in malnourished children has varied over the years, depending on the characteristics of the group of children studied. In 1990, Fischer et al^9^ documented a prevalence of 3% in south West Nigeria. However, other reports gave a higher prevalence in this group of children, ranging from 8.6% in Niger Republic ^10^ to 12% in Central African Republic.^11^ Most of these reports were in severely malnourished children. In 2014, some authors from Northern Nigeria, i,e., Kano, documented a prevalence of 7.8% across the various degrees of malnutrition. These variations were largely attributable to differences in degree of malnutrition and maternal HIV prevalences in the different groups. However, in south western Nigeria there are no current studies to reflect prevalence of HIV status across the spectrum of malnourished children i.e., mild to severe malnutrition. Thus, this study aimed to determine the prevalence and categories of malnutrition amongst children attending two health facilities in Lagos, the Lagos University Teaching Hospital (LUTH) and the Massey Children’s Hospital, and document the HIV status of these malnourished children. In addition, the study aimed to describe any peculiarity in the clinical features, haematological indices and some biochemical indices in these children.

## Materials and Methods

The study was a prospective cross-sectional study conducted at the Paediatric Department of the Lagos University Teaching Hospital and the Massey Street Children’s Hospital over a 6 months period. All children aged 5 months- 5 years diagnosed with malnutrition were consecutively recruited from the emergency room and paediatric wards during the study period. Details of the study and its relevance were explained to parents and caregivers of affected children following which a written, informed consent was obtained.

Known patient with HIV/AIDS on anti-retroviral therapy were excluded from the study.

The sample size was determined with the cochrane formula for prevalence studies. ^12^

The severity of the malnutrition was determined using the using the Weight for Height or z scores (A z-score is the number of standard deviations (SD) below or above the reference median value.) The severity of the Malnutrition was graded as follows: ^13^

1. Moderate malnutrition is weight for height of 70%-80% of expected or z score of between −2SD to < −3SD.
2. Severe malnutrition refers to any of the following:

a. Weight for height <70% or z score of <−3SD, or
b. Mid upper arm circumference <115mm, or
c. Any form of malnutrition with edema

## Data Collection

All the subjects had anthropometry done. The infants were weighed on the bassinet weighing scale (Docbel Braun Model, Doebil Industries, India) while children above 1 year were weighed on a regularly calibrated scale in light clothing. Weight was documented to the nearest Kg. The length of the infants was determined with a standard measuring board (Rolla meter mat 100cm by 1mm) while the height of the older participants was determined to the nearest cm with the stadiometer. The mid upper arm circumference and the Body mass index (BMI) in the children were also determined. BMI (Kg/m2) =Weight in Kg/height in meters^2^.

Pre- and Post- HIV counselling was done by the researchers for all the subjects recruited into the study and all results were treated with confidentiality.

## Laboratory tests

HIV testing was performed using the standard HIV algorithm of two enzyme-linked immunoassays (EIA) in parallel in children above 18 months of age. Real-time polymerase chain reaction (RT-PCR) was performed for children below 18 months of age and those with indeterminate results on EIA. Each subject also had a Full blood count and Serum proteins done.

The following parameters were recorded in a standard proforma for all the subjects:

Age, sex, weight, height/length, presence of oedema, diarrhoea and other significant clinical features, haematological tests (packed cell volume, white blood cell (WBC) count, and differentials, platelet count), HIV tests (ELISA and DNA PCR), Serum Proteins were all documented.

## Statistical Analysis

Data were entered into the SPSS version 21 statistical package and basic descriptive statistics were done. Continuous data and categorical data were compared using the student t test and chi-square test respectively. The factors associated with HIV status were determined with the logistic regression analysis.

## Ethical Considerations

Ethical approval was obtained for the study from the Health Research and Ethics committee of the Lagos University Teaching Hospital. Participants’ privacy and confidentiality of data management was ensured during and after the study.

## Results

### General characteristics of the study population

A total of 214 malnourished children ≤5 years were recruited into the study. 25 of these children were positive for HIV while 189 were HIV negative. Thus, the prevalence of HIV was 11.7% amongst the study participants.

Table 1 shows the general characteristics of the study population in relation to their HIV status. About half of the study participants belonged to the middle socioeconomic class (50.3%) and were Muslims (53.3%).

The median (range) age for the study participants was 15(5-60) months and the median age which was comparable in the HIV positive and negative children. There was no statistical difference in the maternal age (p=0.982), subject’s weight (p=0.052) and length (p=0.086) in the HIV positive and Negative children.

**Table 1:** General characteristics of the study participants.

| **Parameter** | **HIV positive** | **HIV negative** | **P value** |
| --- | --- | --- | --- |
| **Age(months)** |  |  |  |
| Median(range) | 12(9-36) | 12(5-60) |  |
| **Gender (n(%))** |  |  |  |
| Male | 11(44.0) | 83(43.9) | 0.000 |
| Female | 14(56.0) | 106(56.1) |  |
| **Weight** |  |  |  |
| Median(IQR) | 7.75(6.55-8.55) | 7.72(6.00-8.10) | 0.520 |
| **Length/Height** |  |  |  |
| Median(IQR) | 69.75(64.37-75.00) | 70.20(65.00-76.00) | 0.086 |
| **Z score** |  |  |  |
| WHZ | -2.51(-3.50 to -1.37) | -2.50 (-3.83 to -1.50) | 0.224 |
| LHZ | -2.85(-3.45 to -0.95) | -2.85 (-3.63 to -0.460) | 0.606 |
| **Maternal age** |  |  |  |
| Median(IQR) | 29.00(26.00-34.00) | 30.00 (25.00-33.00) | 0.982 |
| **Socioeconomic status N (%)** |  |  |  |
| High | 1(4) | 36(19.0) | 0.162 |
| Middle | 14(56) | 95(50.3) |  |
| Low | 10(40) | 58(30.7) |  |
| **Religion N (%)** |  |  |  |
| Christianity | 12(48) | (37.6) | 0.563 |
| Islam | 12(48) | 112(59.2) |  |
| Others | 1(4) | 16(3.2) |  |

**Median (IQR)= Median(Interquartile range), WHZ-Weight for height z score, LHZ-Length/Height for age z score, n(%)= number(percentage)**

## HIV seropositivity

The prevalence of HIV in the subjects with malnutrition was 11.7%. Most of the HIV positive children were in the age bracket 12 -24 months (60%). However, the highest percentage seropositivity (25.0%) was seen in the age group 25-36 months. The lowest seropositivity was seen in the infants (8.2%) as shown in Table 2.

**Table 2:** Seropositivity of HIV amongst the age categories.

| Variable | HIV Status |  | Seropositivity in each age group |
| --- | --- | --- | --- |
|  | Positive<br>N (%) | Negative<br>N (%) |  |
| Age category (months) |  |  | Percentage (%) |
| <12 months | 5 (20) | 56 (29.6) | 8.2 |
| 12-24 | 15 (60) | 109 (57.7) | 12.1 |
| 25-36 | 3 (12) | 9 (4.8) | 25.0 |
| >36 | 2 (8) | 15 (7.9) | 11.7 |
| Total | 25 (100) | 189(100) | 11.7 |

HIV-Human immunodeficiency virus

With the use of the WHO classification about three quarters of the study participants, had moderate malnutrition (74.8%) while about a quarter (25.2%) had severe forms of malnutrition. (see table 3). 64% of the HIV positive subjects also had moderate malnutrition and almost a quarter of the HIV positive subjects had severe malnutrition. Amongst the HIV positive subjects 16% of the children had severe wasting and 8% had edematous malnutrition.

**Table 3:** Severity of malnutrition and HIV status Severity Classification Z score.

| **Severity Classification**<br>**Z score** | **Total**<br>**N (%)** | **Positive**<br>**N (%)** | **Negative**<br>**N (%)** | **P value** |
| --- | --- | --- | --- | --- |
|  | 214(100) | 25(100) | 189(100) |  |
| -2SD to -3SD | 160(74.8) | 19(76.0) | 141(.74.6) | 0.892 |
| >-3SD | 54(25.2) | 6(24.0) | 48(25.4) | 0.835 |
| Severe wasting | 46(21.5) | 4(16.0) | 42(22.2) | 0.476 |
| Edematous malnutrition | 8(3.7) | 2(8) | 6(3.2) | 0.557 |

**Moderate= z score: between -2SD and -3SD, Severe=z score: >-3SD; p<0.05 is significant, Chisquare (X<sup>2</sup>=3.371), p=0.815**

## Clinical features

Fever, cough and diarrhea were the commonest symptoms seen amongst the study participants while oral thrush, dermatoses and splenomegaly were the commonest signs. (see Table 4)

A higher proportion of the HIV positive subjects compared to the negative subjects had oral thrush, dermatoses and edema but this was not statistically significant. Hepatosplenomegaly was the least presenting signs in both groups. None of the subjects with HIV had otitis media or Jaundice. Splenomegaly was the only significantly different sign among the 2 groups of malnourished children (p=0.0001)

**Table 4:** Clinical characteristics of the malnourished children: HIV positive and HIV negative children.

| **Clinical features** | **HIV Positive**<br>**N=25** | **HIV Negative**<br>**N=189** | **P value** |
| --- | --- | --- | --- |
| Fever | 16(64) | 119(62.9) | 0.919 |
| Cough | 13(52) | 92(48.7) | 0.754 |
| Oral Thrush | 3(12) | 8(4.2) | 0.984 |
| Diarrhea | 5((20) | 56(29.6) | 0.316 |
| Jaundice | 0 | 2(1.1) | 0.605 |
| Edema | 2(8) | 6(3.2) | 0.232 |
| Dermatoses | 3(12) | 17(8.9) | 0.627 |
| Hepatomegaly | 2(8) | 12(6.3) | 0.753 |
| **Splenomegaly** | **2(8)** | **0(0)** | **0.0001** |

HIV-Human immunodeficiency virus

Table 5 shows the laboratory investigations in the study participants. The PCV ranged between 14.1% – 52.5% with a median (IQR) of 35.2 % (28.77-38.10) in the study participants

The median PCV was comparable in both groups. 30(14%) of the study participants had anaemia of which only 2 (0.9%) of the subjects had severe anaemia.

Total WBC ranged from 1,630-56,600cells/mm3 with a median of 11,600/mm3. The WBC and platelets were lower in the HIV positive subjects than the HIV negative subjects but this did not reach statistically significant levels. The proportion of the neutrophil and lymphocytes were also comparable in the two groups.

However, the total protein and serum globulin levels (figure 1) were significantly higher in the HIV positive group (p=0.042, p= 0.015). The serum albumin though lower in the HIV positive group did not reach statistically significant levels. (p= 0.477) CD4 count and CD4% in the HIV group were 645.00 (293.00-1,060) and 12.25 % (10.00-14.25) respectively.

**Table 5:** The laboratory investigations in the study participants.

| **Parameter** | **HIV positive**<br>**Median(IQR)** | **HIV negative**<br>**Median(IQR)** | **P value** |
| --- | --- | --- | --- |
| PCV(%) | 34.75 (28.77-38.10) | 35.40 (32.00-33.00) | 0.391 |
| Total WBC($\times 10^9/L$) | 11.10 (9.4750-15.950) | 11.80 (9.0250-16.175) | 0.603 |
| Neutrophils (%) | 47.40(28.30-62.00) | 47.8 (34.80-59.90) | 0.749 |
| Lymphocytes (%) | 41.40(27.60-57.30) | 41.65 (29.72-54.77) | 0.847 |
| Platelets($\times 10^9/L$) | 431,000(294,000-494,000) | 445,000 (333,000-586,000) | 0.153 |
| **Total protein(g/dl)** | **66.42 (63.11-73.58)** | **63.48 (58.49-68.31)** | **0.042** |
| Serum Albumin(g/dl) | 36.15(28.62-40.82) | 37.15(31.35-41.00) | 0.535 |
| **Serum Globulin** | **30.53(24.61-39.73)** | **27.50 (23.28-31.23)** | **0.015** |

HIV-Human immunodeficiency virus, PCV-Packed cell volume, WBC- White blood cell count. IQR-Interquartile range

The risk factors for HIV evaluated in the study participants are shown in table 7. Multivariate analysis showed that the total protein (p=0.001) and platelet count (p=0.016) were the only factors that could significantly predict the occurrence of HIV in the study population.

**Table 6:** Predictors of the Risk of occurrence of HIV in the malnourished children: Logistic Regression.

| Parameter | B | S.E | Wald | P value | Ex(B)<br>Odd's ratio | 95% Confidence<br>Interval |
| --- | --- | --- | --- | --- | --- | --- |
| Age | 0.327 | 0.318 | 1.055 | 0.304 | 1.386 | 0.743-2.586 |
| Gender | -0.062 | 0.657 | 0.009 | 0.925 | 0.940 | 0.260-3.406 |
| Socioeconomic status | 2.113 | 1.339 | 2.491 | 0.114 | 8.276 | 0.600-114.170 |
| Nutritional status | 1.169 | 0.856 | 1.865 | 0.172 | 3.220 | 0.601-17.242 |
| Edema | -1.575 | 1.324 | 1.416 | 0.234 | 0.207 | 0.015-2.771 |
| Diarrhea | 2.250 | 1.203 | 3.495 | 0.062 | 9.483 | 0.897-100.282 |
| Oral thrush | -0.926 | 1.395 | 0.440 | 0.507 | 0.396 | 0.026-6.107 |
| Fever | -0.193 | 0.635 | 0.093 | 0.761 | 0.824 | 0.237-2.861 |
| **Total Protein** | -0.140 | 0.064 | 4.754 | **0.029** | 0.870 | 0.767-0.986 |
| Globulin | 0.036 | 0.062 | 0.347 | 0.556 | 0.964 | 0.854-1.088 |
| PCV | 0.057 | 0.056 | 1.049 | 0.306 | 1.059 | 0.949-1.180 |
| Total WBC | 0.041 | 0.046 | 0.799 | 0.377 | 1.041 | 0.952-1.140 |
| **Platelet count** | 0.005 | 0.002 | 5.094 | **0.024** | 1.005 | 1.001-1.009 |

P< 0.05 is considered significant, PCV- Packed cell volume, WBC- White blood cell count

## Discussion

In this study, the prevalence of HIV infection amongst malnourished children aged 5 years and below attending two health facilities in Lagos was 11.7%. This prevalence is much higher than the national prevalence for HIV in Nigeria (4.4%) and from other reports in malnourished children in Ogbomosho (3%), ^14^ South Western Nigeria and Kano (7.8%) ^14^ in northern Nigeria. Reports from apparently well children by Ogunbosi et al (10%) ^15^ and Motayo et al (6.7%) have also been observed to be lower.^16^ The possible reasons for the high prevalence observed in this present study may be due to the fact that this was a hospital based study. Other possible explanations may be due to ignorance, weak maternal to child transmission services and failure of early infant diagnosis in the country.

Majority of the subjects in our study were in the age bracket 12-24 months, however the subjects within the age bracket 25-36 months had the highest seropositivity (25%) This age bracket was the peak age of presentation of the infection and probably represented the second phase in HIV infection. It has also been observed that the children who did not receive treatment at this time are likely to die from the illness.^17^ Currently, most children who receive treatment and are compliant now survive into adolescent period and beyond. In terms of sex predilection, there were more females than males in this study which is similar to the findings of Sudawa^17^ and Ogunbosi et al.^15^ Reports from India however document a male preponderance. A female preponderance has also been described in adolescents and adults and this has been attributed to heterosexual transmission. This is not applicable in the paediatric age group and gender differences thus may be a chance finding.^15^

The use of the WHO classification for malnutrition actually reflects the severity of the malnourished state. According to this classification majority of the subjects in this present study had moderate malnutrition and the severe forms was seen in a quarter of the subjects. Edematous malnutrition was present in minority(3.7%) of the subjects. Other reports from south western Nigeria ^18^ have also observed that the less severe forms of malnutrition are no longer prevalent in south western Nigeria.^18^ This may be a reflection of the global effort and efforts at the national level which has resulted in the effective reduction in the severe forms of malnutrition especially in many countries in the sub-Saharan continent.

Majority of the HIV positive subjects in this present study also had moderate malnutrition since this was commonest form of malnutrition seen in the subjects. In contrast, Sudawa et al^17^ in 2013 in Kano state observed that marasmus (48.3%) was the commonest type of malnutrition seen in under fives with HIV infection and only minority (6.5%) had edematous malnutrition. Severe wasting without edema is a common finding in severe malnutrition with concomitant HIV infection.^18–22^ Presently, WHO has corroborated this fact by classifying wasting as seen in Marasmus as a stage 3 AIDS defining illness. Thus, the need arises to always screen children with severe wasting for retroviral disease.

Many studies on malnourished children and HIV status have focused on severely malnourished children as the target group for HIV and the possible reason may be related to the apparent overlap between the two conditions. This present study is one of the few reports which documents the retroviral status across the spectrum of malnourished children in Nigeria and from this present study and other reports it is evident that even children with less severe forms of malnutrition and apparently healthy children are also at risk of the infection.^14–15,17^

In terms of clinical findings, fever, cough and diarrhea were the commonest symptoms seen in the subjects in this present study. These findings are similar to that of previous workers.^17,19,15^ Splenomegaly was seen more in the HIV subjects and appeared to be the only discriminatory clinical feature amongst the HIV positive and negative of malnourished children. Severe malnutrition and HIV have clinical features which overlap and it becomes difficult to differentiate between the two group of patients especially in the resource limited countries however some authors have noted that the severely malnourished HIV infected children were more likely to have oral thrush and persistent diarrhea and these may serve as possible discriminating features.^3,18–19,21–23^

The haematologic indices evaluated in the malnourished children were observed to be lower in the HIV positive group compared with the HIV negative group but this was not statistically significant. This is consistent with the findings of Ezeonwu et al ^24^ in Enugu and Adetifa et al^25^ in Lagos. Lower haematologic indices have been reported in severely malnourished HIV positive children and is probably a reflection of the bone marrow suppression state. Malnourished children are predisposed to anaemia for several reasons namely inadequate intake, micronutrient deficiency (iron and folate deficiencies), increased nutrient loss from recurrent diarrhea. These conditions are prevalent in the HIV infected children and these factors may influence the prevalence of anemia in this group of children.^26^ The aetiology of haematologic abnormalities in the HIV children has been attributed to the release or production of cytokines which occurs in the condition which is believed to suppress haemopoesis and cause myelosuppression.^24^ In addition, other associated comorbid conditions, the presence of opportunistic infections ^27,28^ and the use of antiretroviral drugs^26,30^ especially zidovudine may contribute to the occurrence of anaemia and other haematological abnormalities.^24,29–30^

In contrast, the total protein in the HIV group in this present study was significantly higher than the HIV negative group. A higher level of serum proteins in HIV infection has been reported by other workers^31–33^ and this finding has been attributed to hyperglobulinaemia. The increased total protein levels which occurs in HIV infection is believed to occur as a result of increased immunoglobulins seen in the condition. In our study the globulin level was also found to be higher in the HIV positive subjects compared to the HIV negative children. Theories postulated for the hyperglobulinaemia is the polyclonal B cell activation.^34–36^ The viral proteins (glycoprotein 41) is said to be responsible for the activation of the polyclonal B cell which occurs as the infection progresses.^34–36^ However, the serum albumin was lower in the HIV group compared to the non HIV group which may still be a reflection of the malnutrition state or chronic infection with HIV.

The predictors of the occurrence of HIV in the malnourished subjects were determined with the use of the multivariate analysis and the total protein level and platelet count were able to predict the risk of occurrence of HIV infection in this cohort of malnourished children. Reported predictors/risk factors in malnourished children by Madec et al^10^ however includes gender and previous hospitalizations.

## Limitations

It would have been desirable to follow up the subjects and monitor any changes in the haematological and biochemical profile but this was difficult due to logistic reasons.

## Conclusions

Malnourished children remain a high risk group for HIV infection and the prevalence of the infection obtained in this group of children is still unacceptably high.

The presence of severe wasting or marasmus, oral thrush, diarrhea, splenomegaly, thrombocytopenia and high total proteins in malnourished children should heighten the suspicion of possible underlying associated HIV infection. Continual and mandatory screening of malnourished children for HIV infection is advocated. Further longitudinal studies on malnourished children with HIV is advocated.

## Acknowlegdements

We acknowledge the contribution of the Central research Committee of the University of Lagos for the support for he study. We are also grateful to Mr Niyi and the resident doctors in the department of Paediatrics Lagos University Teaching Hospital who assisted with the recruitment of the subjects and the collection of the data for the study.

## References

1. Fergusson P, Tomskin A, Heikens GT, Bunn J, Amadi B, Manary M, Chagan M et al. Case management of HIV-infected severely malnourished children: challenges in the area of highest prevalence. Lancet 2008; 371: 1305–1307

2. Black RE, Allen LH, Bhutta ZA, Caulfield LE, de Onis M, et al. Maternal and child under nutrition: global and regional exposures and health consequences. Lancet. 2008; 371: 243–260.

3. Bachou H, Tylleskär T, Downing R, Tumwine JK. Severe malnutrition with and without HIV-1 infection in hospitalised children in Kampala, Uganda: differences in clinical features, haematological findings and CD4+ cell counts. Nutrition Journal. 2006; 5:27 doi: 10.1186/1475-2891-5-27

4. Chinkhumba J, Tomkins A, Banda T, Mkangama C, Fergusson P. The impact of HIV on mortality during in-patient rehabilitation of severely malnourished children in Malawi. Trans R Soc Trop Med Hyg. 2008; 102: 639–644.

5. Fergusson P, Tomkins A. HIV prevalence and mortality among children undergoing treatment for severe acute malnutrition in sub-Saharan Africa: a systematic review and meta-analysis. Trans R Soc Trop Med Hyg. 2009; 103: 541–548.

6. Merchant RH, Oswal JS, Bhagwat RV, Karkare J. Clinical profile of HIV infection. Indian Pediatr. 2001; 38: 239–246

7. Kessler L, Daley H, Malenga G, Stephen Graham. The impact of the human immunodeficiency virus type 1 on the management of severe malnutrition in Malawi. Ann Trop Paediatr. 2000; 20:50–56. DOI: 10.1080/02724930092075

8. Van Gend CL, Haadsma ML, Sauer PJ, et al. Delayed Recognition of HIV Infection in malnourished children is associated with poor clinical outcome in low HIV prevalence settings. J Trop Pediatr. 2003; 49:143

9. Fischer GD, Rinaldo CR, Gxbadero D, Kingsley LA, Ndimbie O, Howard C, et al. Seroprevalence of HIV-1 and HIV-2 infection among children diagnosed with protein-calorie malnutrition in Nigeria Epidemiol. Infect. 1993; 110. 373–378

10. Madec Y, Germanaud D, Moya-Alvarez V, Alkassoum W, Issa A, et al. HIV Prevalence and Impact on Renutrition in Children Hospitalised for Severe Malnutrition in Niger: An Argument for More Systematic Screening. PLoS ONE 2011; 6(7): e22787. doi:10.1371/journal.pone.0022787

11. Lesbordes JL, Chassignol S, Manaud F, Salaun D, Bouquety JC, Ray E, et al. Malniutrition and HIV infection in children in the Central African Republic. Lancet 1986: ii: 337–8.

12. Naing L, Winn T, Rush B. Practical issues in calculating the sample size for prevalence studies. Arch Orofacial Sci. 2006; 1:4–9.

13. WHO. Malnurtrition. http://www.who.int/mediacentre/factsheets/malnutrition/en/

14. Adeleke SI, Muktar YM. A Comparative Study of HIV Seroprevalence among Malnourished Children and Controls in Kano. Nigerian Journal of Pharmaceutical Medical Science. 2004; 1: 23–26.

15. Ogubosi BO, Oladokun RE, Brown BJ, Osinusi KI. Prevalence and clinical pattern of pesdiatric HIV infection at the University College Hospital, Ibadan, Nigeria: a prospective cross-sectional study. Ital J Pediatr. 2011; 37: 29.

16. Motayo BO, Usen U, Folarin BO, Okerentugb PO, Innocent-Adiele HC, Okonko IO, Prevalence and Seasonal Variations of HIV 1 and 2 Infection among Children in Abeokuta, South West Nigeria. Journal of Microbiology Research. 2013; 3(3):107–110.

17. Sudawa A, Ahmad AA, Adeleke SA, Umar L, Rogo LD. HIV Infection among Under-Five Malnourished Children in Kano State. World J AIDS. 2013; 3: 350–356

18. Ogunlesi TAl, Ayeni VA, Fetuga BM, Adekanmbi AF. Severe acute malnutrition in a population of hospitalized under-five Nigerian children. Niger Postgrad Med J. 2015; 22(1):15–20.

19. Bugaje MA, Aikhionbare HA. Paediatric HIV/ AIDS seen at Ahmadu Bello University Teaching Hospital Zaria,Nigeria. Ann Afr Med. 2006; 5(2): 73–77.

20. Motayo BO, Usen U, Folarin BO, Okerentugba PO, Innocent-Adiele HC, Okonko IO. Prevalence and Seasonal Variations of HIV 1 and 2 Infection among Children in Abeokuta, South West Nigeria. Journal of Microbiology Research. 2013; 3(3): 107–110.

21. Yeung SD, Wilkinson S, Escott CF, Gilks F. Paediatric HIV Infection in a Rural South Africa District Hospital. J Trop Pediatr. 2000; 46(2):107–110.

22. Babatunde O, Regina EO, Biobele JB, Kikelomo IO. Prevalence and Clinical Pattern of Paediatric HIV Infection at the University College Hospital, Ibadan, Nigeria: A Prospective Cross-Sectional Study. Ital J Pediatr. 2011; 37:1824–7288.

23. Ticklay IM Nathoo KJ, Siziya S, Brady JP: HIV infection in malnourished children in Harare, Zimbabwe. East Afr Med J. 1997; 74: 217–220.

24. Ezeonwu BU, Ikefuna AN, Oguonu T, Okafor HU. Prevalence of hematological abnormalities and malnutrition in HIV-infected under five children in Enugu. Niger J Clin Pract 2014; 17: 303–8.

25. Adetifa IM, Temiye EO, Akinsulie AO, Ezeaka VC, Iroha EO. Haematological abnormalities associated with paediatric HIV/AIDS in Lagos. Ann Trop Paediatr 2006; 26: 121–5

26. Tindyebwa D, Kayita J, Musoke P, Eley B, Nduati R, Coovadia H, Bobart R,Mbori-Ngacha D, Kieffer MP (Eds): Introduction. Handbook on paediatric AIDS in Africa. African network for the care of children affected by AIDS. 2006

27. Evans RH, Scadden DT. Haematological aspects of HIV infection. Baillieres Best Pract Res Clin Haematol 2000;13: 215–30.

28. Okechukwu AA, Gambo D, Okechukwu IO. Prevalence of Anaemia in HIV-Infected Children at the University of Abuja Teaching Hospital, Gwagwalada. Niger J Med 2010; 19:50–7.

29. Johannessen A, Naman E, Gundersen SG, Bruun JN. Antiretroviral treatment reverses HIV-associated anemia in rural Tanzania. BMC Infect Dis 2011; 11: 190. doi.org/10.1186/1471-2334-11-190

30. Ruhinda EN, Bajunirwe F, Kiwanuka J. Anaemia in HIV-infected children: Severity, types and effect on response to HAART. BMC Pediatr 2012; 12:170. doi.org/10.1186/1471-2431-12-170

31. Jemikalajah JD, Adu ME. Assessment of serum proteins in human immunodeficiency virus patients in Auchi, Nigeria. Afr J Cell Pathol 2015; 5: 14–17

32. Ikekpeazu JE, Ibegbu M, Agu NV. Effect of Short and Long Term Exposure of HIV Patients to Highly Active Antiretroviral Therapy (HAART) on Lipid Profile. BAOJ Hiv 2017; 3(1):020

33. Akinpelu OO, Aken’Ova YA andO. Ganiyu Arinola. Levels of Immunoglobulin Classes Are Not Associated with Severity of HIV Infection in Nigerian Patients. World J AIDS. 2012; 2(3): 232–236. doi: 10.4236/wja.2012.23030.

34. Pascale JM, Isaacs MD, Contreras P, Gomez B, Lozano L, Austin E,et al. Immunological markers of disease progression in patients infected with the human immunodeficiency virus. Clin Diagn Lab Immunol. 1997; 4(4):474–477.

35. Arinola OG, Salimonu LS, Okiwelu OH, Muller CP. Levels of Immunoglobulin Classes, Acute Phase Proteins and Serum Electrophoresis in Nigerians Infected with Human Immmunodeficiency Virus. Eur J. Sci. Res. 2005; 7 (3):34–44.

36. Arinola OG, Igbi J. Serum Immunoglobulins and Circulating Immune Complexes in Nigerians with Human Immmunodeficiency Virus and Pulmonary Tuberculosis Infection. Trop J Med Res.1998; 2 (2): 41–48.

